# MATE-Seq: Microfluidic Antigen-TCR Engagement Sequencing

**DOI:** 10.1101/706606

**Authors:** Alphonsus H.C. Ng, Songming Peng, Alexander M. Xu, Won Jun Noh, Katherine Guo, Michael T. Bethune, William Chour, Jongchan Choi, Sung Yang, David Baltimore, James R. Heath

## Abstract

Adaptive immunity is based on peptide antigen recognition. Our ability to harness the immune system for therapeutic gain relies on the discovery of the T cell receptor (TCR) genes that selectively target antigens from infections, mutated proteins, and foreign agents. Here we present a method that selectively labels peptide antigen-specific CD8+ T-cells in human blood using magnetic nanoparticles functionalized with peptide-MHC tetramers, isolates these specific cells within an integrated microfluidic device, and directly amplifies the TCR genes for sequencing. Critically, the identity of the peptide recognized by the TCR is preserved, providing the link between peptide and gene. The platform requires inputs on the order of just 100,000 CD8+ T cells, can be multiplexed for simultaneous analysis of multiple peptides, and performs sorting and isolation on chip. We demonstrate 1000-fold sensitivity enhancement of antigen-specific T-cell receptor detection and simultaneous capture of two virus antigen-specific T-cell receptors from samples of human blood.

## Introduction

A critical aspect of adaptive immunity is the engagement of T-cell receptor (TCR) proteins by agonist antigens presented by major histocompatibility complexes (MHCs). This interaction initiates a cascade of downstream signaling that promotes T cell effector functions such as cytokine secretion.^1–3^ To address the diversity of agonist antigens, the TCR relies on three variable binding domains, called complementarity-determining regions (CDRs). There is an outstanding need to match the TCR genes, especially the sequences encoding CDR3 domains, with their cognate peptide antigen-MHC (pMHC), to better clone therapeutic cells and model this critical interaction. This requires the isolation and TCR sequencing of antigen-specific T cells from a source like peripheral blood mononuclear cells (PBMCs). An intrinsic associated challenge is that any given antigen-specific T cell clone exists at low abundance, and the relatively weak binding affinity of TCRs to their cognate pMHC hampers the selective isolation of an antigen-specific T cell. Furthermore, TCR gene sequencing can also be confounded by the heterodimeric structure of the constituent α and β chains of the TCR and the highly diverse TCR repertoire.

These challenges have prompted the development of approaches that use dye-labeled pMHC tetramers (for increased avidity) and fluorescence activated cell sorting (FACS)^4^ of single cells for TCR analysis. Related methods that use metal ion-labeled^5, 6^ or DNA-labeled^7^ pMHC tetramers have also been reported. Zhang and coworkers^8^ recently reported an approach in which the identity of the antigen is encoded using DNA tags on the pMHCs, thus permitting the linking of TCR sequences to their cognate antigens via single cell sequencing of FACS sorted populations. However, FACS sorting requires a large input of PBMCs, which can be restrictive for the analysis of non-expanded patient derived blood samples. Expansion to generate large enough input can alter population profiles.^9–11^

We recently reported on the use of magnetic iron-oxide nanoparticles (NPs) to support the presentation of >10^4^ pMHC tetramers per particle (Peng et al, *under revision).* These high avidity pMHC-presenting NPs (pNPs), which can also be labeled with multi-functional DNA barcodes, permit an improved efficiency of antigen-specific T cell capture relative to pMHC tetramers. In particular, the pNPs enabled >90% recovery of antigen-specific T cells from a sample input of 10^5^ to 10^6^ CD8+ PBMCs, facilitating the enumeration of antigen-specific T cells directly from unexpanded PBMCs. In principle, these same pNPs could be used as labels for FACS sorting of single antigen-specific T cells for genetic analysis. However, light scattering from the pNP barcodes renders them poor FACS labels, and so we demonstrated TCR sequencing by picking out individual antigen-specific T cells manually. Here we look to develop a higher throughput approach that retains the avidity of pNPs for antigen-specific CD8+ T cell labeling.

We report here on a single-stream Drop-seq derived method,^12^ called microfluidic antigen-TCR engagement sequencing (MATE-seq), for screening of barcoded pNPs against CD8+ T cells. On-chip operations include purifying the pNP-labeled T cells from free pNPs, entrainment of those cells into a droplet-generating microfluidic circuit, and in-drop execution of RT-PCR. The methodology required several advances to seamlessly integrate multiple functions onto a single microfluidic chip, as well as optimization of droplet chemistry for improved stability and efficiency during PCR thermal cycles. We utilized the MATE-seq approach to capture and analyze rare, virus-antigen specific CD8+ T cells from healthy patient bloods.

## Experimental

### Reagents and Materials

Unless otherwise specified, reagents were purchased from Sigma-Aldrich. Deionized water with resistivity of >18 megaohms·cm was used to prepare all aqueous solutions. Roswell Park Memorial Institute (RPMI) 1640 Medium, phosphate buffered saline (PBS), and Calcein AM (C1430) were purchased from ThermoFisher Scientific (Waltham, MA). Fetal bovine serum (FBS) was purchased from ATCC (Mannassas, VA). Penicillin-Streptomycin mixture (17-602E) was purchased from Lonza (Basel, Switzerland). 3-N-Maleimido-6-hydraziniumpyridine hydrochloride (MHPH) and succinimidyl 4-formylbenzoate (SFB) was purchased from TriLink BioTechnologies (San Diego, CA). ImmunoCult™ Human CD3/CD28 T Cell Activator purchased from STEMCELL Technologies Inc. (Vancouver, BC, Canada). Aquapel glass treatment was purchased from Pittsburgh Glass Works LLC (Pittsburgh, PA).

### Production of SAC-DNA conjugates

The oligonucleotide-conjugated streptavidin was produced using cysteine-modified streptavidin (SAC), following a previously published protocol.^13^ Briefly, SAC was first expressed from the pTSA-C plasmid containing the SAC gene (Addgene).^14^ SAC (1 mg/ml) was buffer exchanged to PBS containing Tris(2-Carboxyethyl) phosphine Hydrochloride (TCEP, 5 mM) using Zeba desalting columns (Pierce). Then MHPH (100 mM) in DMF was added to SAC at a molar excess of 300:1. In the meantime, SFB (100mM) in DMF was added to MHC-DNA (500 μM), a 5’-amine modified ssDNA (5’-NH2-AAA AAA AAA A TAG GCA TCC CGA GGA TTC AG), at a 40:1 molar ratio. After reacting at room temperature for 4 hours, MHPH-labeled SAC and SFB-labeled DNA were buffer exchanged to citrate buffer (50 mM sodium citrate, 150 mM NaCl, pH 6.0), and then mixed at a 20:1 (DNA to SAC) molar ratio to react at room temperature overnight. The DNA-SAC conjugate was purified using a Superdex 200 gel filtration column (GE health) and concentrated with 10K MWCO ultra-centrifuge filters (Millipore).

### Production of DNA-labeled peptide-MHC tetramers

We used the method of conditional antigen exchange to enable the rapid construction of HLA-A*02:01 peptide-MHC (pMHC) by the release of a photo-labile peptide, KILGFVFJV.^15–17^ The photo-labile peptide and other peptide antigens were synthesized with standard automated Fmoc-peptide synthesis methodology, where J or (S)-3-(Fmoc-amino)-3-(2-nitrophenyl) propionic acid is the photo-labile amino acid residue. Plasmids encoding human MHC class I heavy chain and human β_2_m containing bacterial strain were kind gifts from Ton N. M. Schumacher. The MHC photo-labile protein (MHC-J) was folded from MHC heavy chain inclusion body, β_2_m inclusion body, and photo-labile peptide according to previously published protocol^18^ and then biotinylated using BirA biotin ligase. A mixture of biotinylated MHC-J (0.5 μM) and peptide antigen (50 μM) was exposed to 365 nm UV light for 45 minutes to generate the biotinylated pMHC. The peptide antigen was either CMV (NLVPMVATV), EBV (GLCTLVAML), MART-1 (ELAGIGILTVI) or NY-ESO (SLLMWITQV) epitope-specific. To form DNA-conjugated pMHC tetramers, SAC-DNA conjugates and biotinylated pMHC were mixed at a 1:4 (streptavidin to MHC) molar ratio, and incubated at room temperature for 30 minutes with rotation.

### Magnetic Nanoparticle Labeling

NPs labeled with pMHC tetramers and DNA primers (pNPs) were formed using streptavidin coated magnetic NPs (500 nm radius, Dynabeads MyOne Streptavidin T1, ThermoFisher catalog #: 65601) according to the manufacturer’s recommended protocol for biotinylated nucleic acid attachment. The NPs were incubated with a mixture of biotinylated DNA (equal parts of PS1-PI-Cα, PS1-PI-Cβ and NP-DNA) at a 1:8 (streptavidin to biotin) molar ratio (Sup. Table 1). The DNA oligomers (synthesized by IDT) were biotinylated at the 5’ end. The biotins of PS1-PI-Cα and PS1-PI-Cβ were attached to the oligomer via a photo-cleavable group (labeled as 5PCBio in Sup. Table 1). The NP-DNA is complimentary to the SAC-DNA. After excess DNA was washed away, the DNA-labeled pMHC tetramers and DNA-labeled NPs were hybridized at 37°C for 30 minutes at a molar ratio of 1.5:1 (MHC-DNA to NP-DNA), and washed once with 0.1% BSA and 2 mM MgCl2 in PBS. Typically, each analysis uses 2.5 μL of stock NPs (28.2 million particles total) per antigen.

### Cell Culture

Frozen leukapheresis fractions from donor NRA11 (donor 1) or NRA13 (donor 3) (UCLA IRB#03-2-023), or commercial peripheral blood mononuclear cells from donor CMV30 (donor 2) (Cellular Technology Limited, Cleveland, OH) were thawed in warm cell culture media (RPMI 1640 containing 10% FBS, 100 U/mL penicillin, and 100 μg/mL streptomycin). Thawed cells were re-suspended in cell culture media supplemented with IL-2 (100 IU/mL) at a density of 10^6^ cells per mL. The cells were first activated for 3 days using CD3/CD28 T Cell activator (1:40 in culture media). After 3 days, cells were resuspended with fresh culture media supplemented with IL-2, expanded in culture for up to 1 week, and frozen until use. Prior to use, the expanded specimens were thawed, and enriched for live CD8+ T cells using a magnetic activated cell sorting kit (Miltenyi Biotech) according to the manufacturer’s instructions.

### Magnetic Isolation of Antigen-specific T cells

Live CD8+ T cells stained with calcein AM (a green-fluorescent live/dead stain, ThermoFisher) were incubated with pNPs for 15 min at room temperature with rotation. Free pNPs and barcoded T cells were enriched by magnetic isolation, and unbound cells were washed away (DynaMag™-2 Magnet, Thermo, 12321D). The free pNPs and barcoded cells were imaged by bright field and epifluorescence microscopy or analyzed by MATE-seq.

### Device Fabrication

The microfluidic device mold was fabricated on silicon wafers using SU-8 2025 (MicroChem). SU-8 was spin coated at 2000 to 4000 rpm for 60 seconds, and photolithography was performed according to the manufacturer’s datasheet to create feature heights ranging from 30 to 55 micrometers, as characterized by a profiler (Dektak). Ultraviolet (UV) light exposure was performed on a Karl Suss MA6 mask aligner, and the features were developed in SU-8 developer (MicroChem). To form the poly(dimethylsiloxane) (PDMS) device, the mold was first treated with chlorotrimethylsilane vapor for 30 minutes, and Sylgard 184 (DOW) comprising 10:1 monomer to crosslinker mixture was poured onto the mold, degassed, and cured at 80°C for 2 hours. The PDMS features were cut out, and 0.7-mm or 2-mm holes were punched through the output or input ports, respectively. PDMS debris was removed using N2 gas, deionized water, and isopropanol. After the PDMS device was dried, dust was removed using tape (Scotch), and the device was plasma-bonded to a glass microscope slide to form the final device.

### Device Preparation

The device has four inputs (I1, I2, I3, and I4) and two outputs (O1 and O2), and comprises two major regions: the deterministic lateral displacement (DLD) size selector and the droplet generator. The droplet generator was made hydrophobic by applying 1 μl of Aquapel to the device through O1. After 30 seconds of incubation, Aquapel was removed, and the device was dried in the oven for 30 minutes at 80°C and cooled to room temperature. Each of the inputs was attached to a reagent reservoir formed from a razor-trimmed 2-200 μL pipette tip, and the O2 output was connected to a 1-mL syringe via 50 cm Tygon tubing (AAD04103-CP ND-100-80, Cole-Parmer). To prime the device for use, four sets of solutions were sequentially infused into the device through O2 while O1 was plugged with a pin: 1) 70% ethanol (to remove air bubbles), 2) 3% F68 pluronic in PBS (to passivate the surface from cell and biomolecular fouling), 3) PBS (to wash out excess pluronic), and 4) buffer (0.73× PBS containing 100 mM NaCl, optimized for in-droplet RT-PCR). Around 800 μl of each solution was infused into the device through the 1-mL syringe at O2 using a syringe pump (New Era Pump Systems, Inc, NE-1000) set at 3 mL/hr. The excess solution, accumulated at the reagent reservoirs, were removed before the next solution was infused.

### Device Operation

After the device was primed, O1 was connected to a 60 mL syringe through 2 meters of Tygon tubing with the syringe plunger positioned at 30 mL. To initiate the droplet generation, HFE-7500 fluorinated oil (3M) with 5% (w/w) PEG-PFPE amphiphilic block copolymer surfactant (008-Fluoro-surfactant, Ran Technologies) was loaded in I3, and a syringe vacuum pressure of around 0.75 atmosphere was applied at O1 by positioning the plunger at 40 mL. The plunger position was held in place by mounting the syringe on a syringe pump (Harvard apparatus, PHD 2000) set at a withdrawal rate of 5 mL/hr. Once droplet generation was confirmed by bright field microscopy, bare pNPs, buffer, and lysis RT-PCR mix (described in the next section) were loaded to I1, I2, and I4, respectively. The O2 withdrawal flow rate (typically between 0.2 to 0.3 mL/hr) was tuned to 1) prevent the NPs from leaking into the droplet generator and 2) maintain a lysis RT-PCR mix to buffer ratio of about 2:1, which was determined visually at the flow focusing junction. Once these conditions were met, bare NPs in I1 were replaced with the sample (i.e. magnetically isolated antigen-specific T cells, comprising free pNPs and barcoded cells). Throughout the sample processing, the oil and lysis RT-PCR mix were replenished as needed, and the buffer level in the I2 reagent reservoir was kept higher or equal to the sample level in the I1 reagent reservoir. The droplets containing lysed cells were collected inside the 2-meter Tygon tubing, which was placed on ice during the device operation to minimize RNA degradation. After the completion of sample processing, pressure was released by temporarily detaching the 60-ml syringe from the Tygon tube. Subsequently, the tubing was disconnected from O1 and placed into a 1.5-mL Eppendorf tube, and the droplets were dispensed into the tube by gently pushing the plunger of the syringe.

### In-Droplet RT-PCR

The lysis RT-PCR mix described above comprises reverse transcriptase, PCR polymerase, dNTPs and buffer from Qiagen one step RT-PCR kit (210212), RNase inhibitor (Promega, 2U/μl), 0.25% IGEPAL CA 630 to lyse the cells, dithiothreitol (DTT, 5mM) to improve RT-PCR efficiency, and multiplexed primers α ID-Vα (Sup. Table 2) and β ID-Vβ (Sup. Table 3). In addition to the polymerase provided in the Qiagen kit, KOD hot DNA polymerase was added (EMD Millipore 71086, 0.02 U/μL) to increase the extension speed and decrease the total time necessary for the RT-PCR process. Excess oil was removed from the collected droplets, and the droplets were distributed in 200 μL PCR tubes. The PCR tubes were placed on ice and exposed to 365nm UV light (UV crosslink cat # 89131-484, VWR) for 10 minutes to release the ssDNA primers (PS1-PI-Cα and PS1-PI-Cβ) from the pNPs. Subsequently, RT-PCR was performed with the following temperature conditions: 50°C for 1 hour, 95°C for 5 min, repeat 40× of (94°C for 10 s, 68°C for 20 s, 70°C for 20 s), 72°C for 10 min, and hold at 12°C. Following the RT-PCR, the amplified DNA products were extracted from the oil droplets by adding an equal volume of 50% perfluorooctanol (PFO) solution in HFE-7500 to the droplets. The droplets were vortexed, centrifuged at 1000g for 2 min, and the supernatant was recovered for subsequent gel purification and PCR.

### Gel Purification, PCR Enrichment, and Sequencing

Following in-droplet RT-PCR, a series of gel purifications and PCR steps were performed to remove interfering oligonucleotides and non-specific amplifications, enrich for the TCR DNA, and prepare the products for sequencing. The amplicons from in-droplet RT-PCR were purified by agarose (1.3%) gel electrophoresis, and the bands were visualized using ethidium bromide. DNA fragments between 400 and 500 base pairs were excised and purified using a Qiagen MinElute Gel Extraction Kit, according to manufacturer’s instructions. The purified DNA was divided in half for enrichment PCR, amplifying TCR α and β gene DNA in separate reactions. The primers used are listed in Sup. Table 4, and PCR was performed with the following conditions: 95°C for 2 min, repeat 30× of (95°C for 20 s, 60°C for 20 s, 70°C for 20 s), 72°C for 10 min, and hold at 12°C. Following enrichment PCR, the amplicons were purified by gel electrophoresis and DNA fragments between 450 and 550 base pairs were excised and extracted, as described above. The purified TCR α and β DNA were prepared for sequencing by adapter insertion PCR. This used primers listed in Sup. Table 5, and PCR was performed with the same conditions as described above, except the primer annealing temperature was 55°C instead of 60°C. Finally, the amplicons were purified by gel electrophoresis and DNA fragments between 500 to 600 base pairs were excised, extracted, and sent out to Laragen (Culver City, CA) for next generation sequencing. The sequencing was carried out on an Illumina Miseq machine using a V2 2 × 150 kit.

### Bulk Sequencing

Around 1000 CD8+ cells were incubated in lysis RT-PCR buffer with RT-PCR primers PS1-PI-Cα, PS1-PI-Cβ, and variable primers α ID-Vα and β ID-Vβ. Samples were processed in an identical fashion to in-droplet samples in preparation for sequencing.

### FACS-isolated Sequencing

Individual cells were sorted into wells of a 96-well plate containing cell lysis buffer (10mM Tris, pH=8, with 1 U/μl RNAse inhibitor, Promega). Following 1 hour lysis at −80 °C, the cell lysates were split in half to perform separate α and β gene amplification and sequenced.^19^

### Sequencing Analysis Pipeline

A computational sequencing analysis pipeline was developed to match reads to peptide-MHC identifiers (PI). The PI of each read was first isolated from the first base pair reads of the paired-end sequencing data (read 1). After sorting reads by PI, DNA sequences were aligned to known TCR genes and processed using MixCR.^20^ For a given antigen, the CDR3 regions of TCR genes were collected. A TCR was considered antigen-specific if >90% of the PIs associated with the TCR were the same. The antigen similarity score of a set of CDR3 sequences was defined as the average Levenshtein distance between each element in the set and the closest element of the entire set of CDR3 sequences that existed in the VDJ database^21^ matched to the antigen. Antigen similarity scores of sequenced CDR3s from MATE-seq experiments were compared to nonspecific similarity scores using 1000 random sets of 12 CDR3 sequences sampled from bulk sequencing of T-cells and CDR3 sequences.

## Results & discussion

The development of MATE-seq (Fig. 1) was motivated by the desire to understand the relationship between specific antigen-presenting MHC complexes and their cognate TCR genes in rare T cell populations. We employed the high avidity of pNPs (Fig. 1A) for antigen-specific T cell capture and TCR α/β gene analysis. The use of these NP reagents required that we address four challenges. First, there are only a handful of cells specific to a particular antigen within a given patient blood draw, and so we needed a microchip platform designed for efficient handling of those rare cell populations (Fig. 1B and Fig. 2). Second, the most efficient way to analyze a specimen is to screen the cells against a library of pNPs, where each library element presents a unique peptide antigen. This required further pNP engineering (primer design of Fig. 1A) so that the antigen identity of a pNP can be linked with the final TCR cDNA library sequences (Fig. 1C). Third, most pNP library elements will not bind to cells, and will need to be removed prior to any sequencing analysis of those cells. Thus, an on-chip purification strategy (Fig. 2A-C) is needed to separate the labeled cells from the unbound pNPs. Finally, labeled cells need to be analyzed individually, which required the incorporation of a Drop-seq type module onto the microchip (droplet generator of Fig 1B, Fig 2D), and the development of Drop-seq chemistries (Fig 1C) to permit reverse transcription and the first round of PCR to be carried out within the microdroplets used to isolate individual cells. We provide, below, detailed descriptions of how each of the challenges were addressed.

**Figure 1:**
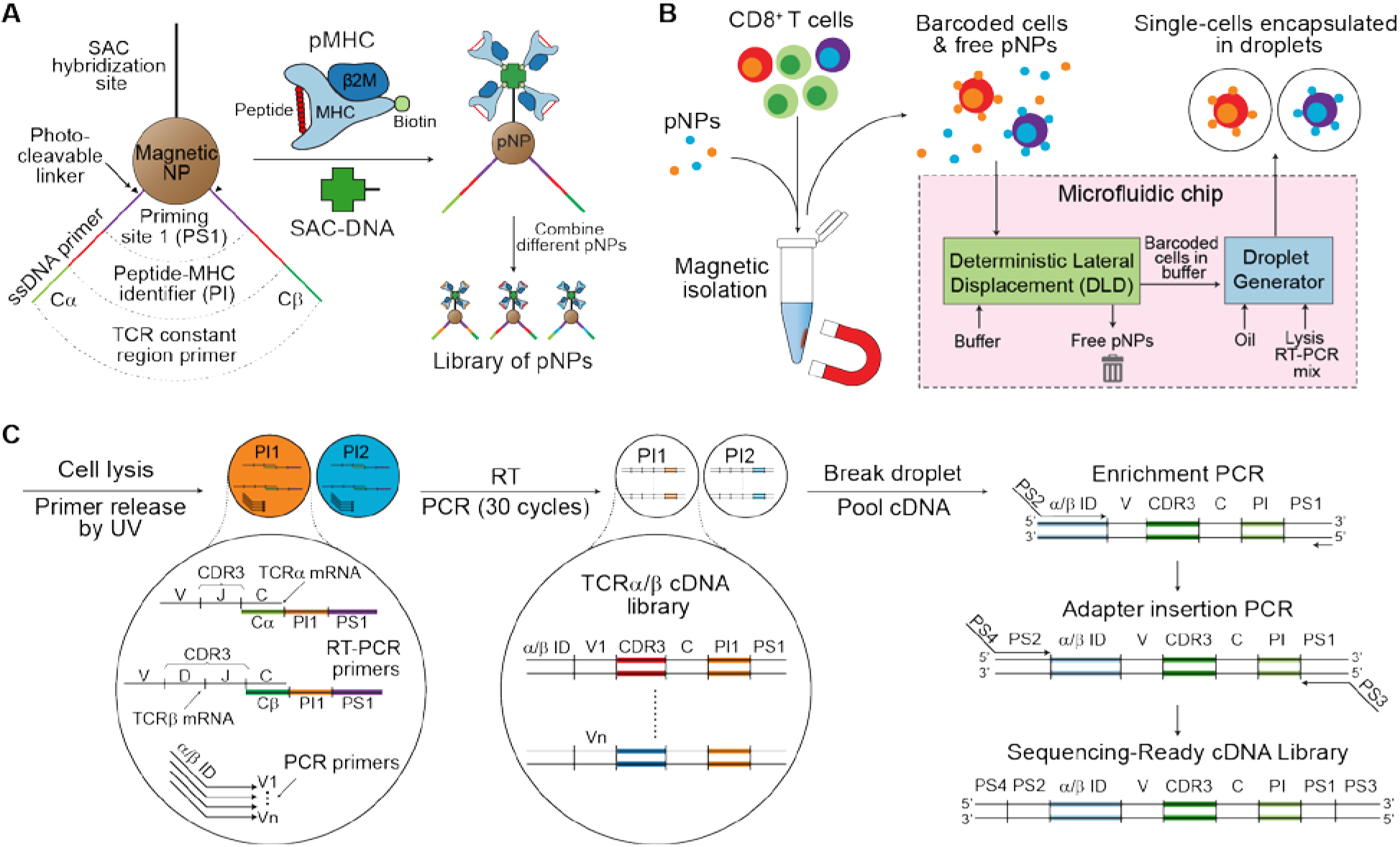
Schematic of MATE-seq. **(A)** MATE-seq relies on magnetic nanoparticles, called pNPs, that present peptide antigens (via tetramerized MHC complexes), as well as photocleavable primers designed to capture TCR α or β chain mRNA for sequencing and transfer the identity of the antigen to the mRNA during RT-PCR. **(B)** A library of antigen-specific pNPs are mixed CD8+ T-cells, and pNP-bound cells (barcoded cells) and unbound pNPs (free pNPs) are purified by magnetic isolation. This mixture is added into a microfluidic device that first removes free pNPs, and then encapsulates barcoded cells in droplets. **(C)** Inside the droplets, individual cells are lysed to release their mRNA, the pNPs are exposed to UV to release the primers, and RT-PCR is performed. The antigen specificity of the pNPs (PI) is transferred to the resulting TCR cDNA library inside the droplet. After droplet breakage, the cDNA libraries are pooled for purification and PCR enrichment, as well as adapter insertion to prepare them for sequencing.

**Figure 2:**
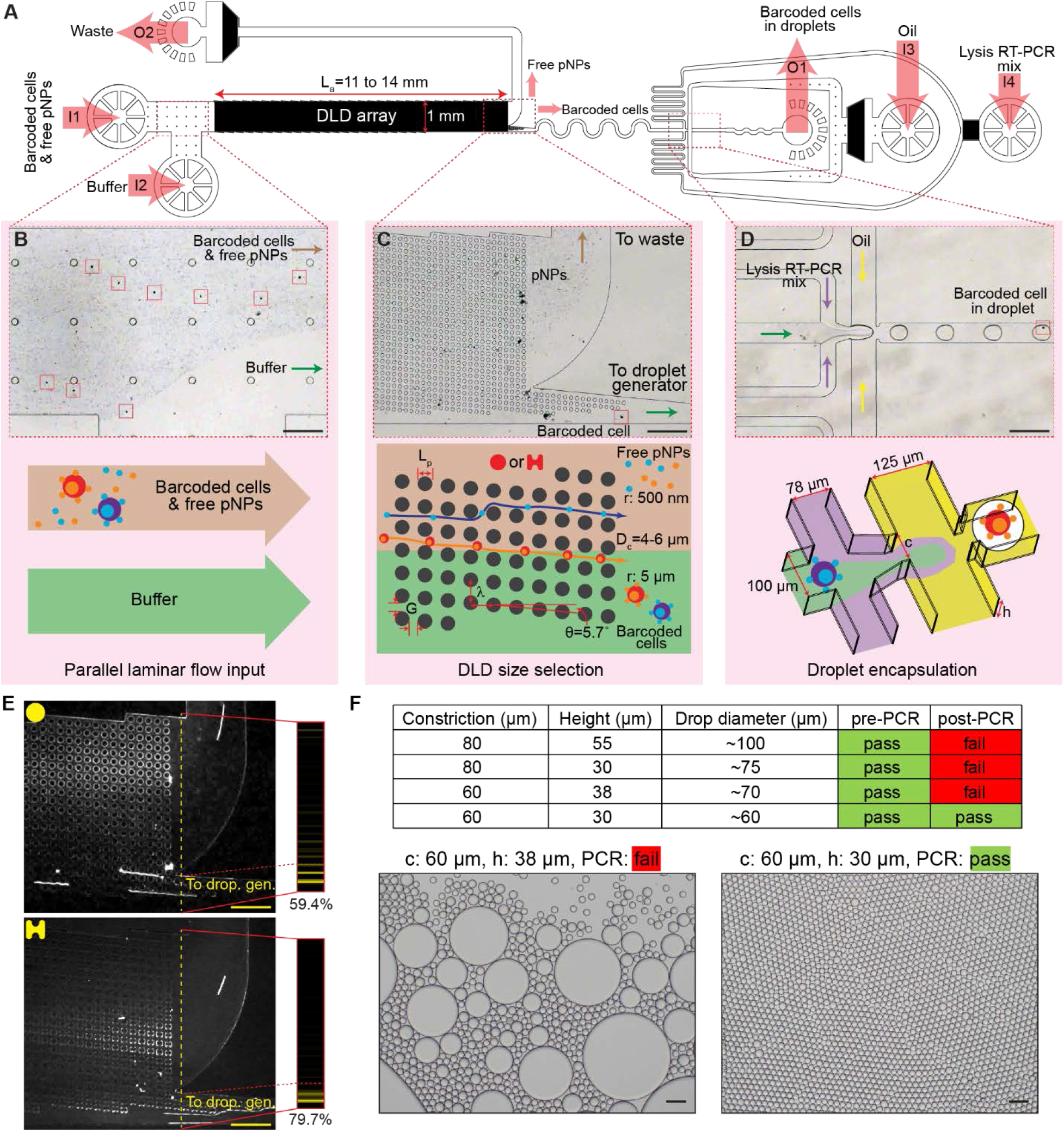
Microfluidic device for MATE-seq. **(A)** The device integrates a DLD array on the front end of a droplet generator, and features 4 inputs (I1 to I4) for barcoded cells-free pNPs mixture, buffer, encapsulation oil, and lysis/RT-PCR mix, as well as 2 outputs (O1 and O2) for the water-in-oil droplets created by the droplet generator and waste. **(B)** The barcoded cells and free pNPs are flowed in parallel with buffer optimized for RT-PCR with minimal mixing by diffusion. **(C)** After DLD processing, the barcoded cells are displaced into the buffer stream toward the droplet generator, with full removal of free pNPs into the waste. The physical dimensions of the DLD array determine a critical sorting diameter, Dc. **(D)** Barcoded cells are encapsulated in water-in-oil droplets with lysis RT-PCR mixture, and collected for analysis. The size of the droplet is determined by the physical dimensions the device. The course of barcoded cells is shown in the micrographs of **A** to **D**, highlighted in solid red squares. **(E)** Fluorescent micrographs of viability-stained cells processed by devices with circle (top) or I-shaped (bottom) pillars. As the cells exit the DLD array, they tend to follow tracks defined by the pillar spacing, as seen in the frequency plot (inset). The intensity of the frequency plot along the y-coordinate is proportional to the number of cells that pass through the device at the yellow dotted line. The I-shaped pillars increased cell sorting efficiency into the droplet generator from ~60% to ~80%. **(F)** The focusing constriction (c) and height (h) of the droplet generator was empirically optimized to produce droplets small enough to withstand 30 cycles of PCR. Droplets with diameter >60 μm appear to merge into larger droplets after PCR, as shown in the bright field micrographs. All scale bars are 200 μm.

### Nanoparticle Design and Validation

To enable TCR sequencing and antigen identification of pNP-labeled cells, we modified the original pNP design (Peng et al, *under revision)* to incorporate TCR-gene specific primers with antigen identifiers, as illustrated in Fig. 1A (not drawn to scale). This new pNP carries three DNA oligomers (Sup. Fig. 1), each performing a specific function. The first oligomer, called NP-DNA, enables the loading (via hybridization) of >10^4^ pMHC tetramers onto each pNP (Peng et al, *under revision).* The second and third oligomers are photo-cleavable primers with three segments for TCR mRNA capture (Cα or Cβ), antigen peptide identification (PI), and RT-PCR priming (PS1). The sequences of these oligomers (NP-DNA, PS1-PI-Cα, and PS1-PI-Cβ) are detailed in Sup. Table 1. These oligomers are attached to the NP via biotin-streptavidin binding, rather than synthesized on-NP as in other bead-supported primer methods,^12^ due to the small size of the NP. The primers enable the generation of a cDNA library containing TCR α and β sequences, each appended with a short PI DNA barcode that identifies the corresponding antigen-specificity (Fig. 1C). In principle, a six-nucleotide PI can encode a library with 4096 elements, although fewer elements are preferable to allow for error correction.^22^ We first validated that particles hybridized with NY-ESO pMHC tetramers can pull down NY-ESO-specific Jurkat cells. As shown in Sup. Fig. 2, all viability stained cells were coated with pNPs, indicating that the pNPs could efficiently enrich the target cells by magnetic pulldown. We found that pNPs with a radius of 500 nm were optimal for magnetic pulldown with minimal particle sedimentation, aggregation, and non-specific cell collection. Most importantly, the small 500-nm radius particles establish a clear size differential from eukaryotic cells, enabling the use of deterministic lateral displacement (DLD)^23^ to separate pNPs from cells, as described in the next section.

### Sample Processing and Microfluidic Filtration

Fig. 1B illustrates the sample processing workflow. First, a library of pNPs (both antigen specific and non-specific) is mixed with CD8+ T cells, and the pNP barcoded cells and free pNPs are magnetically isolated from the non-labeled cells. Next, the barcoded cells and free pNPs are introduced onto the microfluidic platform (Fig. 2A) in parallel with buffer (Fig. 2B), permitting the cells and particles to interact with a DLD array.^23–25^ The array is a periodic arrangement of pillars designed to separate particles larger and smaller than a critical diameter (D_c_). The key parameters and the principle of DLD are shown in Fig. 2C. The array is made of pillars that have a constant center-to-center distance of λ, which is a sum of the pillar length, L_p_, and the gap size, G. Each succeeding column of pillars is vertically shifted down by a distance Δλ with respect to the previous column, and the shift is reset after N columns, which is the period of the DLD array. The period, N, is related to λ and Δλ, and determines the row shift fraction, ε, and displacement angle, θ:

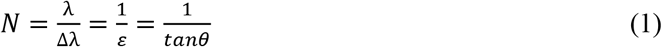

An empirical formula has been previously derived^23^ for the approximation of D_c_ (in μm):

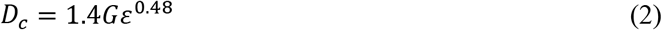

In our DLD design, we set N to 10, and therefore ε = 0.1 and θ = 5.71°. We tested a range of pillar length (L_p_ = 19 to 32.5 μm), gap size (G = 11 to 12.5 μm), center-to-center distance (λ = 30 to 45 μm), and shift distance (Δλ = 3 to 4.5 μm). These parameters result in a critical diameter for sorting, D_c_, of around 4 to 6 μm, meaning that any cell above this size range will be separated from free pNPs. We also tested different array lengths (L_a_ = 11 to 14 mm) and shapes (I-post or circles),^23–28^ which have been shown to affect sorting performance. As illustrated in Fig. 2C, the free pNPs (radius of 500 nm, <D_c_) follow a “zigzag” trajectory toward the waste, while the barcoded cells (radius > 5 μm, >D_c_) are displaced into the buffer stream toward the droplet generator (Fig. 2C).

In the optimized designs (L_a_ = 14 mm, G = 12.5 μm, λ = 45 μm, and Δλ = 4.5 μm), using either circular or I-shaped pillars, we achieve 100% removal of free pNPs (Fig. 2B and Sup. Movie 1) with a cell isolation efficiency between 60-80% (Sup. Movie 2 and 3), as represented by kymographs of flow (Sup. Fig. 3) and the flow density plot of viability-stained cells at the point of separation (Fig. 2E). This is critical to prevent non-specific PI barcodes from being amplified during in-droplet RT-PCR, as discussed in the next section.

### In-Droplet RT-PCR Optimization

Droplet encapsulation of barcoded cells gives us the opportunity to link TCR genes with the specific peptide antigen bound to the T cell (Fig. 1B). Here, we performed RT-PCR directly in droplets, which maximizes capture of TCR mRNA templates and limits crosstalk of PI barcodes between cells (Fig. 1C). To amplify the TCR genes in droplets, we had to address several challenges. First, it was critical to maximize the thermal stability of the droplets so that they can withstand at least 30 cycles of PCR. We included up to 5% w/w PEG-PFPE amphiphilic block copolymer surfactant in the oil phase to stabilize the droplet interface, but still observed that larger droplets tended to burst during thermal cycling. However, if the droplet was too small, there were usually insufficient reagents to complete the RT-PCR reaction and low quality libraries were generated. The size of these droplets is determined by the nozzle height (h), flow focusing constriction (c), and flow rates of the droplet generator (Fig. 2D). We determined empirically that the maximum droplet size with the required stability was 60 μm in diameter, which was generated using a device that had a flow focusing nozzle height and constriction of 30 and 60 μm, respectively (Fig. 2F).

Another challenge was tuning the protocol and reagent design of the in-droplet RT-PCR. Inside each droplet, a cell is lysed and its mRNA contents are released, enabling the priming of the TCR mRNA by PS1-PI-Cα or PS1-PI-Cβ, released from the pNPs via photocleavage (Fig 1C). RT-PCR directly on the pNPs without releasing the primer exhibited poor efficiency. After reverse transcription, the resulting cDNA is amplified using multiplexed primers α ID-Vα (Sup. Table 2) and β ID-Vβ (Sup. Table 3), which are included in the lysis RT-PCR reagent to prime the variable segments of the TCR. The primers bind close to the junction of the constant region and variable region of the TCR so that 150 base reads are sufficient to cover the CDR3 region. The in-droplet PCR appends the peptide identifier and flanking priming sites (PS1 and α ID or β ID) to the TCR cDNA, yielding the final product in the droplet: α/β ID-V-CDR3-C-PI-PS1 (Fig. 1C).

### Device Integration

The MATE-seq device and workflow were designed to accommodate several experimental conditions. First, the input sample volume is low—typically <50 microliters of barcoded cell/pNP mixture is obtained from a pulldown, often with only a handful of barcoded cells. Such low sample volumes preclude syringe injection due to the high dead volume of the syringe, adapter, and tubing. Further, direct coupling of DLD separation to droplet generation requires a coordination of flow rates that can be difficult to achieve in practice. Conventional droplet generation microchips utilize syringe pumps, with pump flow rates precisely optimized. For example, the oil inlet is typically pumped two to three times faster than the aqueous inlets (e.g. cell or RT-PCR reagents).^12^ However, in the MATE-seq device, such a strategy would require coordination of 4 syringe pumps. We devised a simpler approach in which two syringe pumps were utilized to withdraw fluid at the outputs while, for the inputs, we used pipette tips as low-volume input reagent reservoirs (see Methods).

In summary, operation of the MATE-seq device required effective separation and purification of free pNPs from barcoded cells, followed by cell isolation and RT-PCR within droplets. The pNPs played two roles. First, they were used to selectively bind T-cells using TCR-pMHC specific interactions. Second, they transferred a peptide identifier (PI) barcode so that, after in-droplet RT-PCR, the TCR α and β genes from a specific cell carried the same PI barcode. The optimized process flow permitted efficient operation without additional sorting equipment. After microfluidic processing, the RT-PCR product undergoes two more bulk PCR and gel purification steps prior to sequencing (Fig. 3A) (see Methods). An optimized microfluidic design is supplied as a CAD file here (Supplementary Materials).

**Figure 3:**
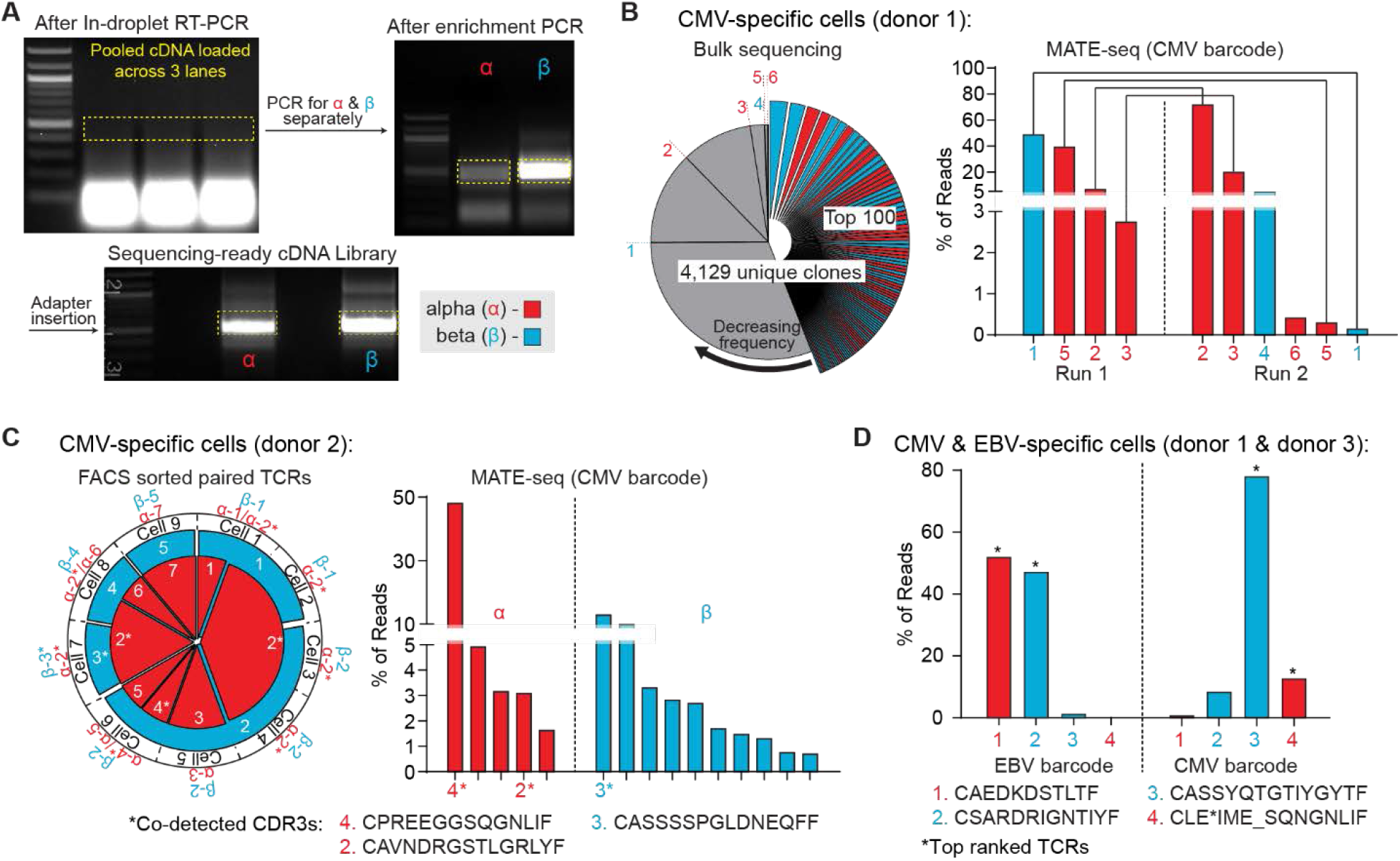
MATE-seq sequencing results. **(A)** After microfluidic processing, the RT-PCR product was purified and modified with two PCR steps to generate a sequencing-ready library. Gel ladder is 100 base pairs. **(B)** The bulk analysis of cells containing CMV-specific cells (donor 1) yielded 4,129 sequenced TCR clones, with no single TCR contributing more than 3% of total reads. Two independent MATE-seq analyses of the same cells yielded 2 unique TCR β chains and 4 unique TCR α chains, all of which were only associated with CMV barcodes, with 1 β and 3 α chains shared across both runs. While these 6 CDR3s contributed to >99% of total reads in MATE-seq, they only represented 0.09% of total reads from bulk sequencing. **(C)** FACS sorting and sequencing of a different set of CMV-specific T-cells (donor 2) yield 9 paired TCRs, with the most common α and β chains appearing in 6 and 4 cells, respectively. MATE-seq analysis of the same cells co-detected 3 of the same CDR3s, one of which was the most common α chain from FACS analysis. **(D)** Multiplexed MATE-seq of CMV and EBV-specific cells (mixture of donor 1 and 3) revealed dominant TCR α and β chains for each antigen.

### Comparison to Bulk Analysis

We compared a bulk sequencing approach with MATE-seq in the analysis of PBMCs with known CMV-specific minorities (donor 1). Performing bulk sequencing on ~1000 of the PBMCs yielded a background of 144,635 unique TCR gene sequences comprising 4,129 CDR3 clones, with the most frequent TCR clone (α or β) consisting of 2.4 percent of the total reads (Fig. 3B, left). In the MATE-seq analysis, we screened the specimen against a library of pNPs, including one EBV and one CMV HLA-A*02:01 antigen, MHC-J (as a control), and the MART-1 antigen, each with a unique PI barcode. In two independent analyses, 300-500 barcoded cells were processed by the microchip device, yielding 2 unique TCR β chains and 4 unique TCR α chains with 1 β and 3 α chains shared across both runs (Fig. 3B, right, Sup. Table 6). These 6 TCR genes, which were associated with the CMV barcode, represented 99.97 and 99.56 percent of total reads detected after MATE-seq enrichment runs, respectively. In contrast, the same CMV-specific TCRs represented the 400^th^, 726^th^, 1387^th^, 1870^th^, 1918^th^, and 2869^th^ most common clones from the bulk TCR analysis, occupying 0.09% of total reads. Thus, MATE-seq enriched CMV-specific genes by approximately 10^3^ times, and the PI barcode provided a level of error correction against non-specific pNP pulldowns.

### Comparison to Standard FACS Method

We compared a standard FACS method with MATE-seq in the analysis of another set of PBMCs with known CMV specificity (donor 2). We FACS sorted and sequenced 9 single cells, yielding paired TCRs with 7 unique α and 5 unique β chains, as well as 4 instances of 2 α chains being isolated from the same single cells (Fig. 3C, left, Sup. Table 7).^29, 30^ Using MATE-seq, we processed ~300 cells on a microfluidic device and sequenced their TCR genes. This resulted in a total of 5 α and 10 β chains, including 2 of the 3 α chains and 1 of the 6 β chains obtained from FACS sorting (Fig. 3C, right, Sup. Table 8). These results demonstrate that MATE-seq was able to isolate some of the same TCR genes at high-throughput as the gold standard FACS method.^19^

### Multiplexed Analysis

Single cell encapsulation by the droplet generator allows MATE-seq to perform multiplexed TCR sequencing while maintaining the identity of the antigen bound in the TCR-MHC complex for each cell. We demonstrated this capability in the analysis of PBMCs from a mixture of donors with known CMV and EBV specificity (mixture of donor 1 and donor 3) (Fig. 3D). After sequencing, only 4 unique TCR genes were detected above the background (>99.8% of reads). Pairing the reads to the EBV or CMV barcodes showed that each epitope was paired with a dominant TCR pair, with good alignment with database predictions, as described below. We did find some evidence of cross-talk, likely due to barcode switching occurring after droplet breakage. For example, the 3^rd^ and 4^th^ ranked EBV TCRs were the 1^st^ and 2^nd^ ranked CMV TCRs. Thus, we only consider the top ranked TCRs for each antigen.

### Public Database Comparison

The antigen specificity of a sequenced TCR gene can be supported through comparison against public databases. For example, at the time of analysis, there were some 6,689 CMV (NLVPMVATV) antigen-specific TCR CDR3s reported^21^. TCRs identified against this antigen using MATE-seq should show similarities to this database that are not found with random TCR sequences.

We defined the antigen similarity score of a CDR3 region as the minimum number of additions, deletions, or substitutions needed to convert a sequenced CDR3 to an antigen-specific CDR3 database element. An exact match, for example, yields a score of 0 (Fig. 4A).^21^ Subsets of randomly sampled CDR3s from bulk sequencing (1000 subsamples of 12 CDR3s) yield an average EBV similarity score of ~5.2 ±0.6 and an average CMV similarity score of ~4.1±0.5. By contrast, antigens identified by MATE-seq as CMV-specific yield an average CMV similarity score of 2 (donor 2, n=13), 3 (donor 1, n=6), and 3 (donor 1 multiplexed analysis, n=2), and EBV-specific antigens yield an average EBV similarity score of 0. 5 (donor 3 multiplexed analysis, n=2). All combined CMV specific cells from the MATE-seq runs yield a CMV similarity score of 2.38 (n=21), while FACS sorted cells yield an average CMV similarity score of 2.17 (donor 2, n=12).

**Figure 4:**
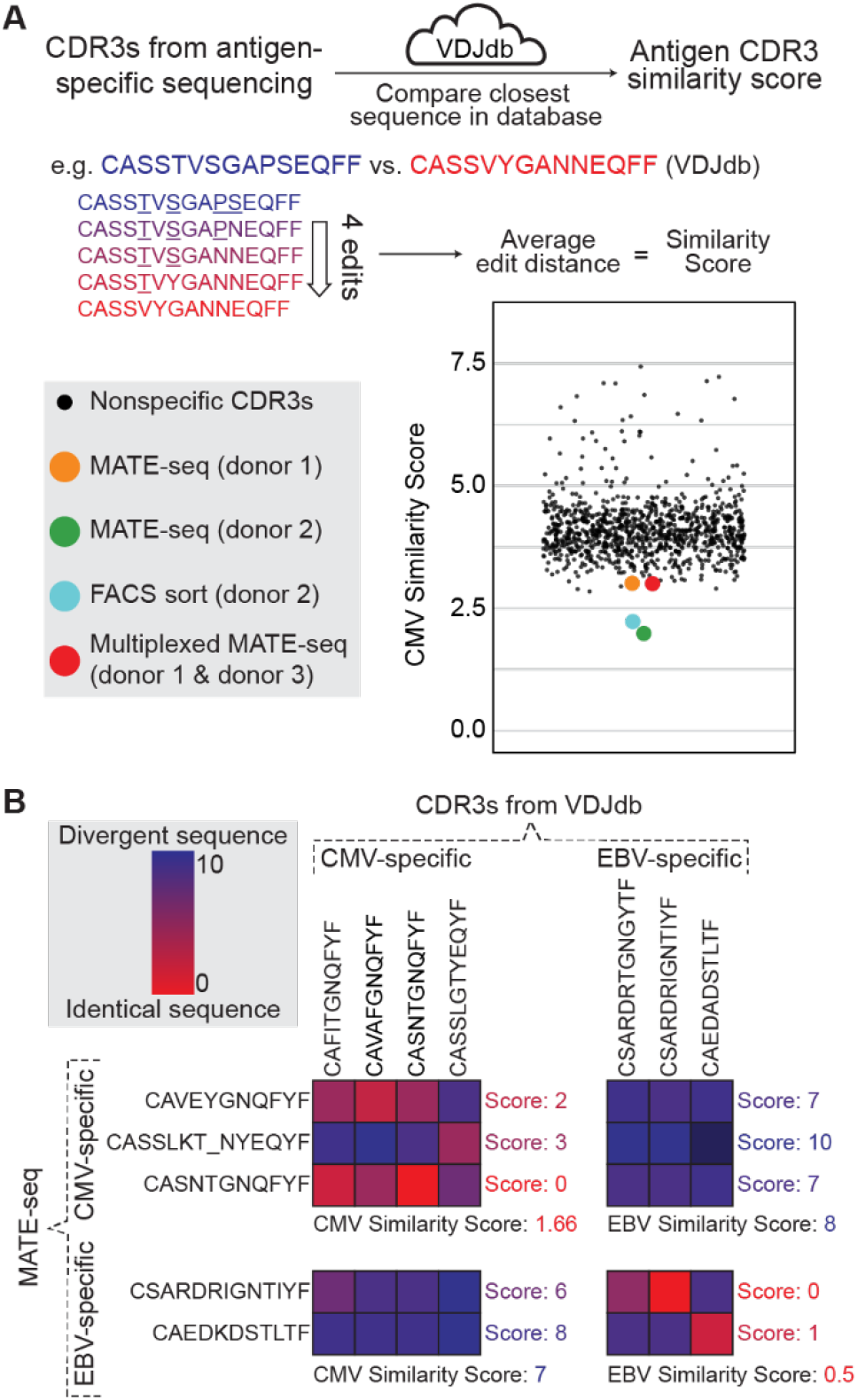
Comparison of MATE-seq with public TCR database. **(A)** Sequenced TCR CDR3 regions were compared to a public database of reported antigen-specific CDR3s. The antigen similarity score for an antigen-specific sequencing was defined as the average edit distance of each sequenced CDR3 to its nearest antigen-specific CDR3 in the database. **(B)** Example of CMV and EBV antigen similarity score matrix. Cross-antigen similarity scores showed that MATE-seq derived CDR3s for CMV and EBV were likely antigen-specific and far more similar to database CDR3s for the same antigen. In particular, MATE-seq uncovered one EBV and three CMV CDR3s with exact match to the database (i.e. score: 0) – one exact match from each antigen is shown.

Exact matches between sequenced TCR genes and database TCR genes for specific antigen were found, but were rare. In fact, of the TCR genes reported in the database specific to CMV cells, only 8 CDR3 genes were reported 10 times or more out of 7,186 total entries. Of these total entries 6,689 were unique, demonstrating the vast reported diversity of TCR genes and explaining the relative infrequency of exact CDR3 region matches. Yet, MATE-seq analysis uncovered exact matches for three CMV-specific CDR3 (CASNTGNQFYF, CAWSVSDLAKNIQYF, CASSYQTGTIYGYTF) and one EBV-specific CDR3 (CSARDRIGNTIYF) (Fig. 4B). These results demonstrate the capacity of MATE-seq to isolate likely antigen-specific T-cells for sequencing.

## Conclusion

The MATE-seq method utilizes the high avidity of MHC tetramer-loaded magnetic NPs to enable sensitive capture of antigen specific T cell populations. Additional engineering modifications of the NP scaffold, plus a custom designed microfluidic chip, enabled further genetic analysis of individual captured T cells, so that the TCR α and β gene sequences from a given T cell are matched with the antigen specificity of that cell. The full process relied upon 3 technology innovations. First, the NPs provided a modular platform to streamline multiplexing by mixing combinations of primers and binding reagents on the same substrate. Each pNP concentrates >10^4^ MHC tetramers to increase the avidity of the typically weak TCR-pMHC interaction beyond what is achieved using pMHC tetramers.^31^ Second, the seamless integration of DLD and droplet encapsulation enabled individual purified pNP-labeled T cells to be captured into droplets using relatively simple and robust operating parameters. Finally, droplet chemistry optimizations allowed for the first round of PCR to be carried out within the microdroplets, while retaining droplet integrity.

Here we demonstrate this technology by capturing and sequencing a couple of virus antigen-specific T cell populations from healthy patient blood, with limited multiplexing. Future work will focus on increasing the library size of screened pMHCs, and on improving the chemistry associated with the indrop RT-PCR steps, so as to increase the information extracted from each cell.^12, 32^ For example, paired TCR α and β chain sequences can also be generated using bridging primers, to directly link a cell’s α and β TCR genes physically before sequencing.^33^ By adding gene-specific primers for relevant T-cell genes^34^ or generic polyT primers^12, 32^ to the pNPs, cellular phenotyping can also be carried out so that, for example, individual cells can be probed for signatures of antigen experience, exhaustion, or functional capacity.

## Author contributions

AHCN, SP, AMX, WJN, KG, MTB, JC, SY, DB, and JRH designed the experiments. AHCN, SP, WJN, KG, WC did the experiments. AHCN, SP, AMX, WJN and JRH analyzed the data. AHCN, AMX, and JRH wrote the paper.

## Acknowledgements

Sequencing data is available at the Sequence Read Archive under project PRJNA546025. AHCN is supported by a Banting Postdoctoral Fellowship from the Government of Canada. AMX is supported by a Ruth L. Kirschstein F32 Postdoctoral Fellowship from the National Cancer Institute (1F32CA213966-01). This work was supported by the Parker Institute for Cancer Immunotherapy, by a Caltech-GIST project grant, by the Washington State CARE Foundation, and by the National Cancer Institute (grant #U54CA199090, PI Heath). We thank Drs. Jesse Zaretsky and Antoni Ribas for providing donor specimens.

