## Supplementary Figs. and Tables for "MATE-Seq: Microfluidic Antigen-TCR Engagement Sequencing"

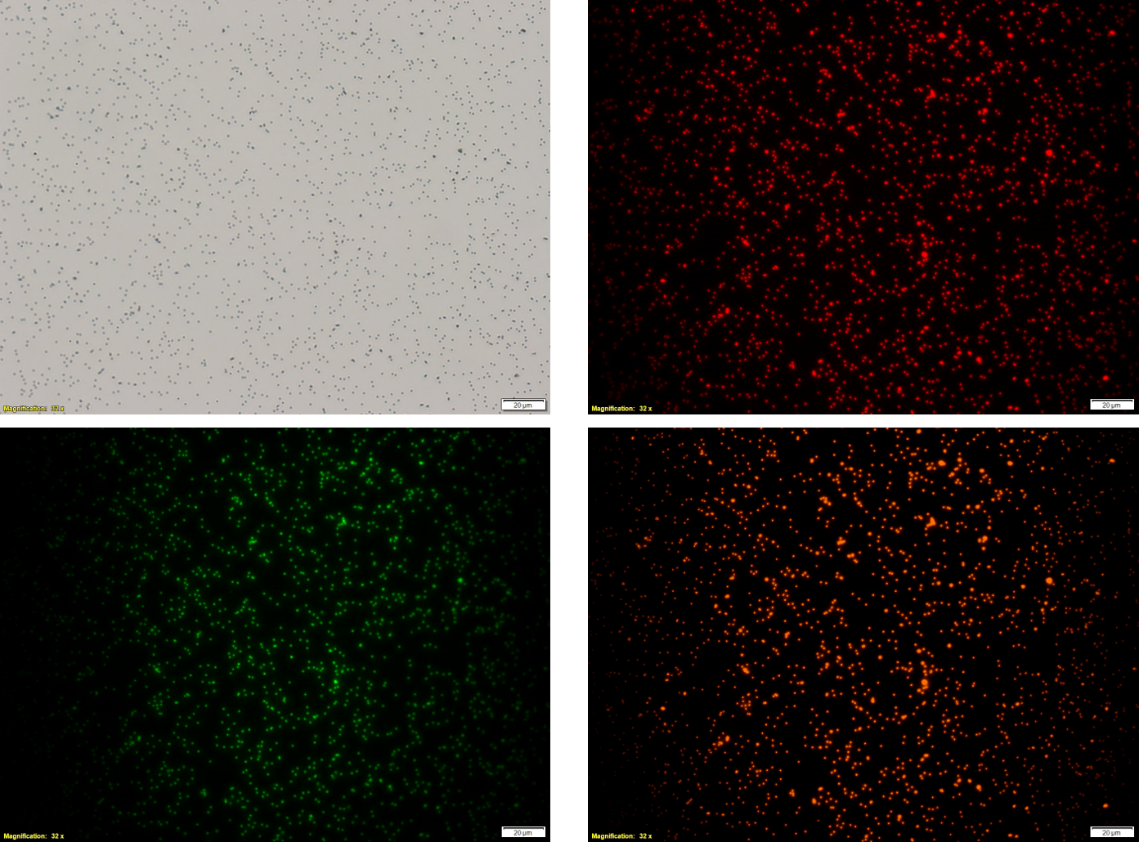

**Supplementary Figure 1:** Bright field and fluorescent micrographs of pNPs. The pNPs are labeled with three different DNA (PS1-PI-Cα, PS1-PI-Cβ, NP-DNA) at the same time. Complementary DNA for Cα (Red), Cβ (Green) and MHC-DNA (Yellow) are hybridized to the DNA on the pNP and give the correct fluorescent emission, indicating successful DNA labeling on the particles.

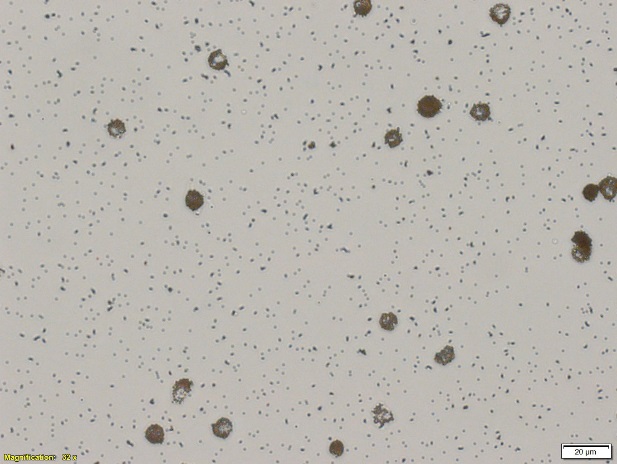

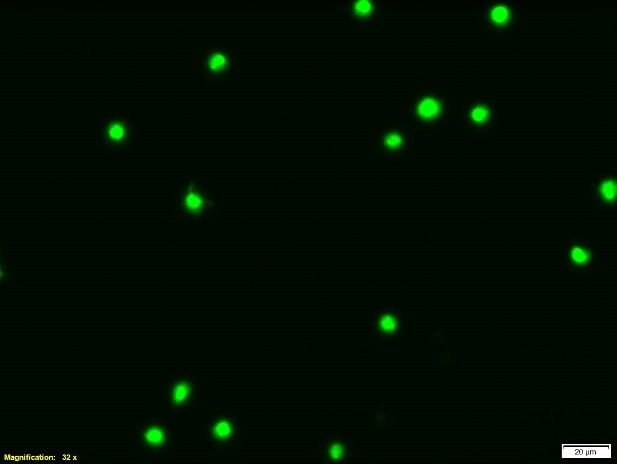

**Supplementary Figure 2**: Bright field and fluorescent micrographs of viability-stained NY-ESO-specific Jurkat cells pulldown by NY-ESO-specific pNPs.

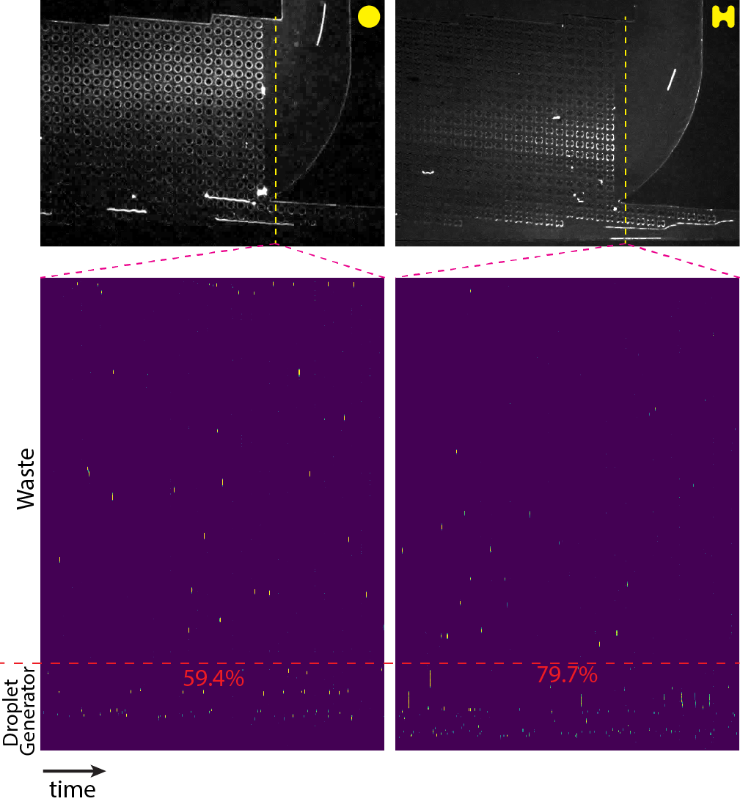

**Supplementary Figure 3:** A movie screen capture of DLD sorting of viability-stained cells using a device with circular (left) or I-shaped pillars (right). As a cell passes the dotted yellow line in the device, the event is recorded as a spot on the corresponding kymograph, below. The fraction of cells that go to the droplet generator is the sorting efficiency of DLD.

**Supplementary Table 1:** DNA sequences used for NP labeling

| **Name** | **DNA sequence** |
| --- | --- |
| PS1-PI-Cα | /5PCBio/TCGTCGGCAGCGTCAGATGTGTATAAGAGACAGNNNNNNNN**XXXXXX**TCTCTCAGCTGGTACACGGC |
| PS1-PI-Cβ | /5PCBio/TCGTCGGCAGCGTCAGATGTGTATAAGAGACAGNNNNNNNN**XXXXXX**GATGGCTCAAACACAGCGACCTC |
| NP-DNA | /5BiosG/AAAAAAAAACTATGTCGATACAAGTCAGATAGTTCAACTCGTTCACTATAA**CTGAATCCTCGGGATGCCTA** |
| N is either one of A, T, G or C  **XXXXXX** is either one of ACTCTT, CGAGTC, TAGACG or TTCAGG, representing peptide identifiers for CMV, MHC-J, MART-1, or EBV, respectively.  **CTGAATCCTCGGGATGCCTA** is the sequence that hybridizes to SAC-DNA | |

**Supplementary Table 2:** Vα-gene-specific primers for cloning TCRα genes

| **TRAV gene** | **α ID** | **Vα** |
| --- | --- | --- |
| TRAV1-1*01 | 5'-TACAGGAAGCCTCAGCA | GGACAAAGCCTTGAGCAGCCCTC-3' |
| TRAV1-2*01 | 5'-TACAGGAAGCCTCAGCA | GGACAAAACATTGACCAGCCCACTG-3' |
| TRAV2*01 | 5'-TACAGGAAGCCTCAGCA | AAGGACCAAGTGTTTCAGCCTTCCAC-3' |
| TRAV3*01 | 5'-TACAGGAAGCCTCAGCA | GCTCAGTCAGTGGCTCAGCCGGA-3' |
| TRAV4*01 | 5'-TACAGGAAGCCTCAGCA | CTTGCTAAGACCACCCAGCCCATC-3' |
| TRAV5*01 | 5'-TACAGGAAGCCTCAGCA | GGAGAGGATGTGGAGCAGAGTCTTTTCC-3' |
| TRAV6*01 | 5'-TACAGGAAGCCTCAGCA | AGCCAAAAGATAGAACAGAATTCCGAGGC-3' |
| TRAV6*03 | 5'-TACAGGAAGCCTCAGCA | GAGGCCCTGAACATTCAGGAGGG-3' |
| TRAV7*01 | 5'-TACAGGAAGCCTCAGCA | GAAAACCAGGTGGAGCACAGCCC-3' |
| TRAV8-1*01 | 5'-TACAGGAAGCCTCAGCA | GCCCAGTCTGTGAGCCAGCATAACC-3' |
| TRAV8-2*01 | 5'-TACAGGAAGCCTCAGCA | GCCCAGTCGGTGACCCAGCTTG-3' |
| TRAV8-2*02 | 5'-TACAGGAAGCCTCAGCA | GCCCAGTCGGTGACCCAGCTTAG-3' |
| TRAV8-3*01 | 5'-TACAGGAAGCCTCAGCA | GCCCAGTCAGTGACCCAGCCTG-3' |
| TRAV8-4*06 | 5'-TACAGGAAGCCTCAGCA | CTCTTCTGGTATGTGCAATACCCCAACC-3' |
| TRAV8-4*07 | 5'-TACAGGAAGCCTCAGCA | GTTGAACCATATCTCTTCTGGTATGTGCAATACC-3' |
| TRAV8-6*01 | 5'-TACAGGAAGCCTCAGCA | GCCCAGTCTGTGACCCAGCTTGAC-3' |
| TRAV8-7*01 | 5'-TACAGGAAGCCTCAGCA | ACCCAGTCGGTGACCCAGCTTG-3' |
| TRAV9-1*01 | 5'-TACAGGAAGCCTCAGCA | GGAGATTCAGTGGTCCAGACAGAAGGC-3' |
| TRAV9-2*01 | 5'-TACAGGAAGCCTCAGCA | GGAAATTCAGTGACCCAGATGGAAGG-3' |
| TRAV9-2*02 | 5'-TACAGGAAGCCTCAGCA | GGAGATTCAGTGACCCAGATGGAAGG-3' |
| TRAV10*01 | 5'-TACAGGAAGCCTCAGCA | AAAAACCAAGTGGAGCAGAGTCCTCAGTC-3' |
| TRAV11*01 | 5'-TACAGGAAGCCTCAGCA | CTACATACACTGGAGCAGAGTCCTTCATTCC-3' |
| TRAV12-1*01 | 5'-TACAGGAAGCCTCAGCA | CGGAAGGAGGTGGAGCAGGATCC-3' |
| TRAV12-2*01 | 5'-TACAGGAAGCCTCAGCA | CAGAAGGAGGTGGAGCAGAATTCTGG-3' |
| TRAV12-2*03 | 5'-TACAGGAAGCCTCAGCA | GGACCCCTCAGTGTTCCAGAGGG-3' |
| TRAV12-3*01 | 5'-TACAGGAAGCCTCAGCA | CAGAAGGAGGTGGAGCAGGATCCTG-3' |
| TRAV13-1*02 | 5'-TACAGGAAGCCTCAGCA | GGAGAGAATGTGGAGCAGCATCCTTC-3' |
| TRAV13-2*01 | 5'-TACAGGAAGCCTCAGCA | GGAGAGAGTGTGGGGCTGCATCTTC-3' |
| TRAV14/DV4*01 | 5'-TACAGGAAGCCTCAGCA | GCCCAGAAGATAACTCAAACCCAACCAG-3' |
| TRAV14/DV4*04 | 5'-TACAGGAAGCCTCAGCA | CAGAAGATAACTCAAACCCAACCAGGAATG-3' |
| TRAV16*01 | 5'-TACAGGAAGCCTCAGCA | GCCCAGAGAGTGACTCAGCCCGA-3' |
| TRAV17*01 | 5'-TACAGGAAGCCTCAGCA | AGTCAACAGGGAGAAGAGGATCCTCAGG-3' |
| TRAV18*01 | 5'-TACAGGAAGCCTCAGCA | GGAGACTCGGTTACCCAGACAGAAGG-3' |
| TRAV19*01 | 5'-TACAGGAAGCCTCAGCA | GCTCAGAAGGTAACTCAAGCGCAGACTG-3' |
| TRAV20*01 | 5'-TACAGGAAGCCTCAGCA | GAAGACCAGGTGACGCAGAGTCCC-3' |
| TRAV21*01 | 5'-TACAGGAAGCCTCAGCA | AAACAGGAGGTGACGCAGATTCCTGC-3' |
| TRAV22*01 | 5'-TACAGGAAGCCTCAGCA | GGAATACAAGTGGAGCAGAGTCCTCCAG-3' |
| TRAV23/DV6*01 | 5'-TACAGGAAGCCTCAGCA | CAGCAGCAGGTGAAACAAAGTCCTCA-3' |
| TRAV23/DV6*04 | 5'-TACAGGAAGCCTCAGCA | CAGCAGGTGAAACAAAGTCCTCAATCTTTG-3' |
| TRAV24*01 | 5'-TACAGGAAGCCTCAGCA | ATACTGAACGTGGAACAAAGTCCTCAGTCAC-3' |
| TRAV25*01 | 5'-TACAGGAAGCCTCAGCA | GGACAACAGGTAATGCAAATTCCTCAGTACC-3' |
| TRAV26-1*01 | 5'-TACAGGAAGCCTCAGCA | GATGCTAAGACCACCCAGCCCCC-3' |
| TRAV26-1*02 | 5'-TACAGGAAGCCTCAGCA | GATGCTAAGACCACCCAGCCCACC-3' |
| TRAV26-2*01 | 5'-TACAGGAAGCCTCAGCA | GATGCTAAGACCACACAGCCAAATTCAATG-3' |
| TRAV27*01 | 5'-TACAGGAAGCCTCAGCA | ACCCAGCTGCTGGAGCAGAGCC-3' |
| TRAV29/DV5*01 | 5'-TACAGGAAGCCTCAGCA | GACCAGCAAGTTAAGCAAAATTCACCATC-3' |
| TRAV30*01 | 5'-TACAGGAAGCCTCAGCA | CAACAACCAGTGCAGAGTCCTCAAGC-3' |
| TRAV34*01 | 5'-TACAGGAAGCCTCAGCA | AGCCAAGAACTGGAGCAGAGTCCTCAG-3' |
| TRAV35*01 | 5'-TACAGGAAGCCTCAGCA | GGTCAACAGCTGAATCAGAGTCCTCAATC-3' |
| TRAV36/DV7*01 | 5'-TACAGGAAGCCTCAGCA | GAAGACAAGGTGGTACAAAGCCCTCTATCTC-3' |
| TRAV36/DV7*02 | 5'-TACAGGAAGCCTCAGCA | GAAGACAAGGTGGTACAAAGCCCTCAATC-3' |
| TRAV38-1*01 | 5'-TACAGGAAGCCTCAGCA | GCCCAGACAGTCACTCAGTCTCAACCAG-3' |
| TRAV38-1*04 | 5'-TACAGGAAGCCTCAGCA | GCCCAGACAGTCACTCAGTCCCAGC-3' |
| TRAV38-2/DV8*01 | 5'-TACAGGAAGCCTCAGCA | GCTCAGACAGTCACTCAGTCTCAACCAGAG-3' |
| TRAV39*01 | 5'-TACAGGAAGCCTCAGCA | GAGCTGAAAGTGGAACAAAACCCTCTGTTC-3' |
| TRAV40*01 | 5'-TACAGGAAGCCTCAGCA | AGCAATTCAGTCAAGCAGACGGGC-3' |
| TRAV41*01 | 5'-TACAGGAAGCCTCAGCA | AAAAATGAAGTGGAGCAGAGTCCTCAGAAC-3' |

**Supplementary Table 3:** Vβ-gene-specific primers for cloning TCRβ genes

| **TRBV gene** | **β ID** | **Vβ** |
| --- | --- | --- |
| TRBV1*01 | 5'-CAGGAGGGCTCGGCA | GATACTGGAATTACCCAGACACCAAAATACCTG-3' |
| TRBV2*01 | 5'-CAGGAGGGCTCGGCA | GAACCTGAAGTCACCCAGACTCCCAG-3' |
| TRBV3-1*01 | 5'-CAGGAGGGCTCGGCA | GACACAGCTGTTTCCCAGACTCCAAAATAC-3' |
| TRBV3-2*01 | 5'-CAGGAGGGCTCGGCA | GACACAGCCGTTTCCCAGACTCCA-3' |
| TRBV4-1*01 | 5'-CAGGAGGGCTCGGCA | GACACTGAAGTTACCCAGACACCAAAACAC-3' |
| TRBV4-1*02 | 5'-CAGGAGGGCTCGGCA | CACCTGGTCATGGGAATGACAAATAAGAAG-3' |
| TRBV4-2*01 | 5'-CAGGAGGGCTCGGCA | GAAACGGGAGTTACGCAGACACCAAG-3' |
| TRBV4-3*04 | 5'-CAGGAGGGCTCGGCA | AAGAAGTCTTTGAAATGTGAACAACATCTGGG-3' |
| TRBV5-1*01 | 5'-CAGGAGGGCTCGGCA | AAGGCTGGAGTCACTCAAACTCCAAGATATC-3' |
| TRBV5-1*02 | 5'-CAGGAGGGCTCGGCA | AGGGCTGGGGTCACTCAAACTCC-3' |
| TRBV5-3*01 | 5'-CAGGAGGGCTCGGCA | GAGGCTGGAGTCACCCAAAGTCCC-3' |
| TRBV5-4*01 | 5'-CAGGAGGGCTCGGCA | GAGACTGGAGTCACCCAAAGTCCCAC-3' |
| TRBV5-4*03 | 5'-CAGGAGGGCTCGGCA | CAGCAAGTGACACTGAGATGCTCTTCTCAG-3' |
| TRBV5-4*04 | 5'-CAGGAGGGCTCGGCA | ACTGTGTCCTGGTACCAACAGGCCCT-3' |
| TRBV5-5*01 | 5'-CAGGAGGGCTCGGCA | GACGCTGGAGTCACCCAAAGTCC-3' |
| TRBV5-8*01 | 5'-CAGGAGGGCTCGGCA | GAGGCTGGAGTCACACAAAGTCCCAC-3' |
| TRBV5-8*02 | 5'-CAGGAGGGCTCGGCA | AGGACAGCAAGCGACTCTGAGATGC-3' |
| TRBV6-1*01 | 5'-CAGGAGGGCTCGGCA | AATGCTGGTGTCACTCAGACCCCA-3' |
| TRBV6-4*01 | 5'-CAGGAGGGCTCGGCA | ATTGCTGGGATCACCCAGGCAC-3' |
| TRBV6-4*02 | 5'-CAGGAGGGCTCGGCA | ACTGCTGGGATCACCCAGGCAC-3' |
| TRBV7-1*01 | 5'-CAGGAGGGCTCGGCA | GGTGCTGGAGTCTCCCAGTCCCTG-3' |
| TRBV7-2*01 | 5'-CAGGAGGGCTCGGCA | GGAGCTGGAGTCTCCCAGTCCCC-3' |
| TRBV7-2*04 | 5'-CAGGAGGGCTCGGCA | GGAGCTGGAGTTTCCCAGTCCCC-3' |
| TRBV7-3*01 | 5'-CAGGAGGGCTCGGCA | GGTGCTGGAGTCTCCCAGACCC-3' |
| TRBV7-3*05 | 5'-CAGGAGGGCTCGGCA | TGGGAGCTCAGGTGTGATCCAATTTC-3' |
| TRBV7-4*01 | 5'-CAGGAGGGCTCGGCA | GGTGCTGGAGTCTCCCAGTCCC-3' |
| TRBV7-6*01 | 5'-CAGGAGGGCTCGGCA | GGTGCTGGAGTCTCCCAGTCTCCC-3' |
| TRBV7-9*01 | 5'-CAGGAGGGCTCGGCA | GATACTGGAGTCTCCCAGAACCCCAG-3' |
| TRBV7-9*03 | 5'-CAGGAGGGCTCGGCA | GATACTGGAGTCTCCCAGGACCCCAG-3' |
| TRBV7-9*04 | 5'-CAGGAGGGCTCGGCA | ATATCTGGAGTCTCCCACAACCCCAGAC-3' |
| TRBV7-9*07 | 5'-CAGGAGGGCTCGGCA | CACAACCGCCTTTATTGGTACCGACAG-3' |
| TRBV9*01 | 5'-CAGGAGGGCTCGGCA | GATTCTGGAGTCACACAAACCCCAAAGC-3' |
| TRBV10-1*01 | 5'-CAGGAGGGCTCGGCA | GATGCTGAAATCACCCAGAGCCCAAG-3' |
| TRBV10-2*01 | 5'-CAGGAGGGCTCGGCA | GATGCTGGAATCACCCAGAGCCCA-3' |
| TRBV10-2*02 | 5'-CAGGAGGGCTCGGCA | AAGGCAGGTGACCTTGATGTGTCACC-3' |
| TRBV11-1*01 | 5'-CAGGAGGGCTCGGCA | GAAGCTGAAGTTGCCCAGTCCCC-3' |
| TRBV11-2*01 | 5'-CAGGAGGGCTCGGCA | GAAGCTGGAGTTGCCCAGTCTCCCAG-3' |
| TRBV11-3*01 | 5'-CAGGAGGGCTCGGCA | GAAGCTGGAGTGGTTCAGTCTCCCAGA-3' |
| TRBV11-3*03 | 5'-CAGGAGGGCTCGGCA | GGTCTCCCAGATATAAGATTATAGAGAAGAAACAGC-3' |
| TRBV12-1*01 | 5'-CAGGAGGGCTCGGCA | GATGCTGGTGTTATCCAGTCACCCAGG-3' |
| TRBV12-2*01 | 5'-CAGGAGGGCTCGGCA | GATGCTGGCATTATCCAGTCACCCAAG-3' |
| TRBV12-3*01 | 5'-CAGGAGGGCTCGGCA | GATGCTGGAGTTATCCAGTCACCCC-3' |
| TRBV12-5*01 | 5'-CAGGAGGGCTCGGCA | GATGCTAGAGTCACCCAGACACCAAGG-3' |
| TRBV13*01 | 5'-CAGGAGGGCTCGGCA | GCTGCTGGAGTCATCCAGTCCCC-3' |
| TRBV14*01 | 5'-CAGGAGGGCTCGGCA | GAAGCTGGAGTTACTCAGTTCCCCAGC-3' |
| TRBV15*01 | 5'-CAGGAGGGCTCGGCA | GATGCCATGGTCATCCAGAACCCAAG-3' |
| TRBV16*01 | 5'-CAGGAGGGCTCGGCA | GGTGAAGAAGTCGCCCAGACTCCA-3' |
| TRBV17*01 | 5'-CAGGAGGGCTCGGCA | GAGCCTGGAGTCAGCCAGACCC-3' |
| TRBV18*01 | 5'-CAGGAGGGCTCGGCA | AATGCCGGCGTCATGCAGAAC-3' |
| TRBV19*01 | 5'-CAGGAGGGCTCGGCA | GATGGTGGAATCACTCAGTCCCCAAAG-3' |
| TRBV20-1*01 | 5'-CAGGAGGGCTCGGCA | GGTGCTGTCGTCTCTCAACATCCGAG-3' |
| TRBV20/OR9-2*01 | 5'-CAGGAGGGCTCGGCA | AGTGCTGTCGTCTCTCAACATCCGAG-3' |
| TRBV21-1*01 | 5'-CAGGAGGGCTCGGCA | GACACCAAGGTCACCCAGAGACCTAGAC-3' |
| TRBV21/OR9-2*01 | 5'-CAGGAGGGCTCGGCA | GACACCAAGGTCACCCAGAGACCTAGATTTC-3' |
| TRBV23-1*01 | 5'-CAGGAGGGCTCGGCA | CATGCCAAAGTCACACAGACTCCAGG-3' |
| TRBV24-1*01 | 5'-CAGGAGGGCTCGGCA | GATGCTGATGTTACCCAGACCCCAAG-3' |
| TRBV25-1*01 | 5'-CAGGAGGGCTCGGCA | GAAGCTGACATCTACCAGACCCCAAGATAC-3' |
| TRBV26*01 | 5'-CAGGAGGGCTCGGCA | GATGCTGTAGTTACACAATTCCCAAGACACAG-3' |
| TRBV26/OR9-2*01 | 5'-CAGGAGGGCTCGGCA | GATGCTGTAGTTACACAATTCTCAAGACACAGAATC-3' |
| TRBV27*01 | 5'-CAGGAGGGCTCGGCA | GAAGCCCAAGTGACCCAGAACCC-3' |
| TRBV28*01 | 5'-CAGGAGGGCTCGGCA | GATGTGAAAGTAACCCAGAGCTCGAGATATC-3' |
| TRBV29-1*01 | 5'-CAGGAGGGCTCGGCA | AGTGCTGTCATCTCTCAAAAGCCAAGC-3' |
| TRBV29-1*03 | 5'-CAGGAGGGCTCGGCA | ACGATCCAGTGTCAAGTCGATAGCCAAG-3' |
| TRBV30*01 | 5'-CAGGAGGGCTCGGCA | TCTCAGACTATTCATCAATGGCCAGCG-3' |
| TRBV30*04 | 5'-CAGGAGGGCTCGGCA | ACTATTCATCAATGGCCAGCGACCC-3' |

**Supplementary Table 4:** Primers for Enrichment PCR

| **Name** | **DNA Sequence** |
| --- | --- |
| PS1 partial | 5’-TCGTCGGCAGCGTCAGATG-3’ |
| PS2-α ID | 5’-GTCTCGTGGGCTCGGAGATGTGTATAAGAGACAGTACAGGAAGCCTCAGCA-3’ |
| PS2-β ID | 5’-GTCTCGTGGGCTCGGAGATGTGTATAAGAGACAGCAGGAGGGCTCGGCA-3’ |

**Supplementary Table 5.** Primers for Adapter Insertion PCR

| **Name** | **DNA Sequence** |
| --- | --- |
| PS3-PS1 | 5’-AATGATACGGCGACCACCGAGATCTACAC[i5]TCGTCGGCAGCGTCAGATGTGTATAAGAGACAG-3’ |
| PS4-PS2 | 5’-CAAGCAGAAGACGGCATACGAGAT[i7]GTCTCGTGGGCTCGGAGATGTGTATAAGAGACAG-3’ |

**Supplementary Table 6.** Donor 1 TCR α and β genes and CDR3 amino acid sequences.

|  | **CDR3** | **Gene** |
| --- | --- | --- |
| **Run 1** | CASSTVSGAPSEQFF | TRBV6-5*00 |
|  | CALYFFFGNEKLTF | TRAV16*00 |
|  | CLE*IME_SQGNLIF | TRAV4*00 |
|  | CAVEYGNQFYF | TRAV3*00 |
| **Run 2** | CLE*IME_SQGNLIF | TRAV4*00 |
|  | CAVEYGNQFYF | TRAV3*00 |
|  | CASSLKT_NYEQYF | TRBV12-4*00 |
|  | CASNTGNQFYF | TRAV24*00 |
|  | CALYFFFGNEKLTF | TRAV16*00 |
|  | CLE*IME_S*GNLIF | TRBV6-5*00 |

**Supplementary Table 7.** Donor 2 TCR α and β genes and CDR3 amino acid sequences from FACS.

|  | **α CDR3** | **Gene** |  | **β CDR3** | **Gene** |
| --- | --- | --- | --- | --- | --- |
| α-1 | CAFMTSG~ATNKLIF | TRAV38-1 | β-1 | CSVWSFGDRDGYTF | TRBV29-1 |
| α-2 | CAVNDRGSTLGRLYF | TRAV12-2 | β-2 | CASSSANYGYTF | TRBV12-3 |
| α-3 | CAVNFGGGKLIF | TRAV12-2 | β-3 | CASSSSPGLDNEQFF | TRBV7-6 |
| α-4 | CPREEGGSQGNLIF | TRAV26-1 | β-4 | CASSYITGTGSYGYTF | TRBV6-5 |
| α-5 | CAAAVETSGSRLTF | TRAV21 | β-5 | CSARDRIGNTIYF | TRBV20-1 |
| α-6 | CARNTGNQFYF | TRAV24 |  |  |  |
| α-7 | CTSPNTNAGKSTF | TRAV26-1 |  |  |  |

**Supplementary Table 8.** Donor 2 TCR α and β genes and CDR3 amino acid sequences from MATE-seq.

| **α CDR3** | **Gene** | **FACS?** | **β CDR3** | **Gene** | **FACS?** |
| --- | --- | --- | --- | --- | --- |
| CPREEGGSQGNLIF | TRAV26-1 | α-4 | CASSSSPGLDNEQFF | TRBV7-6 | β-3 |
| CKASKIIF | TRAV3 |  | CASSEGSWGSDTQYF | TRBV6-5 |  |
| CAVRGDYKLSF | TRAV1-2 |  | RSVQDQDGFTEETQYF | TRBV29-1 |  |
| CAVNDRGSTLGRLYF | TRAV12-2 | α-2 | CASSLGAGPYNEQFF | TRBV18 |  |
| CIAQNNNDMRF | TRAV26-2 |  | CASSGPISSYNEQFF | TRBV6-1 |  |
|  |  |  | CAWSVSDLAKNIQYF | TRBV30 |  |
|  |  |  | CASSLSFGTEAFF | TRBV6-4 |  |
|  |  |  | CASSYPGTGIHGYTF | TRBV6-5 |  |
|  |  |  | CASSLGPSSYEQYF | TRBV5-1 |  |
|  |  |  | CASSEQAGGYGYTF | TRBV6-1 |  |

**Supplementary Movie 1** shows the removal of free pNP to waste and displacement of barcoded cells to the droplet generator, where individual cells are encapsulated in water-in-oil droplets with lysis RT-PCR mix.

**Supplementary Movie 2** shows a fluorescent sequence of viability-stained cells at the output of a DLD array with circular posts.

**Supplementary Movie 3** shows a fluorescent sequence of viability-stained cells at the output of a DLD array with I-posts.
